# Estimating the dimensionality of the manifold underlying multi-electrode neural recordings

**DOI:** 10.1101/2020.12.17.423196

**Authors:** Ege Altan, Sara A. Solla, Lee E. Miller, Eric J. Perreault

## Abstract

It is generally accepted that the number of neurons in a given brain area far exceeds the information that area encodes. For example, motor areas of the human brain contain tens of millions of neurons that control the activation of tens or at most hundreds of muscles. This massive redundancy implies the covariation of many neurons, which constrains the population activity to a low-dimensional manifold within the space of all possible patterns of neural activity. To gain a conceptual understanding of the complexity of the neural activity within a manifold, it is useful to estimate its dimensionality, which quantifies the number of degrees of freedom required to describe the observed population activity without significant information loss. While there are many algorithms for dimensionality estimation, we do not know which are well suited for analyzing neural activity. The objective of this study was to evaluate the efficacy of several representative algorithms for estimating linearly and nonlinearly embedded data. We generated synthetic neural recordings with known intrinsic dimensionality and used them to test the algorithms’ accuracy and robustness. We emulated some of the important challenges associated with experimental data by adding noise, altering the nature of the embedding from the low-dimensional manifold to the high-dimensional recordings, varying the dimensionality of the manifold, and limiting the amount of available data. We demonstrated that linear algorithms overestimate the dimensionality of nonlinear, noise-free data. In cases of high noise, most algorithms overestimated dimensionality. We thus developed a denoising algorithm based on deep learning, the “Joint Autoencoder”, which significantly improved subsequent dimensionality estimation. Critically, we found that all algorithms failed when the dimensionality was high (above 20) or when the amount of data used for estimation was low. Based on the challenges we observed, we formulated a pipeline for estimating the dimensionality of experimental neural data.

**Author Summary:** The number of neurons that we can record from has increased exponentially for decades; today we can simultaneously record from thousands of neurons. However, the individual firing rates are highly redundant. One approach to identifying important features from redundant data is to estimate the dimensionality of the neural recordings, which represents the number of degrees of freedom required to describe the data without significant information loss. Better understanding of dimensionality may also uncover the mechanisms of computation within a neural circuit. Circuits carrying out complex computations might be higher-dimensional than those carrying out simpler computations. Typically, studies have quantified neural dimensionality using one of several available methods despite a lack of consensus on which method would be most appropriate for neural data. In this work, we used several methods to investigate the accuracy of simulated neural data with properties mimicking those of actual neural recordings. Based on these results, we devised an analysis pipeline to estimate the dimensionality of neural recordings. Our work will allow scientists to extract informative features from a large number of highly redundant neurons, as well as quantify the complexity of information encoded by these neurons.

## Introduction

Studies that simultaneously record the activity of many neurons have shown that cortical neural activity is highly redundant [1]. In primary motor cortex (M1), redundancy arises as tens of millions of neurons control tens or at most hundreds of muscles. This redundancy implies significant covariation in the activity of many neurons, which confines the population neural activity to a low-dimensional manifold embedded in the neural space of all possible patterns of neural population activity [2-9]. Low-dimensional manifolds have also been observed in a variety of other cortical regions [10-18]. Reliable algorithms for identifying these manifolds and characterizing their dimensionality are increasingly important as our ability to record from large populations of neurons increases [19]. The dimensionality of the manifold describing the coordinated firing of a set of neurons quantifies the number of degrees of freedom needed to describe population activity without significant information loss [20, 21]. Projecting the observed firing patterns onto the manifold yields a low-dimensional set of latent signals that can simplify the interpretation of population neural activity [2, 9, 22]. Low-dimensional latent signals can facilitate the manipulation or the extraction of signals for brain-computer interfaces, a rehabilitative technology that converts neural signals into control commands to restore movement to paralyzed patients [23, 24].

Unfortunately, it is surprisingly difficult to estimate the dimensionality of neural manifolds, particularly in the realistic condition of a noisy, nonlinear embedding. There is evidence of a nonlinear mapping between the recorded neural activity and the associated low-dimensional latent signals [10, 25-27]. Noise propagates from the level of sensory transduction and amplification, the opening and closing of voltage-gated ion channels, and builds up at the level of synapses, causing neural firing to be a stochastic process [28]. The two effects, nonlinearity and noise, combine to pose significant challenges to existing dimensionality estimation algorithms. The accuracy of the estimators also depends on the amount of available data [29, 30], which is limited in most experimental paradigms. If we wish to identify the manifolds associated with experimentally measured neural activity, we need methods that are robust in the presence of these challenges.

The methods that have been proposed for estimating the dimensionality of neural manifolds can be broadly categorized into linear or nonlinear algorithms, based on assumptions about the nature of the mapping between the low-dimensional representation of the latent signals and the high-dimensional space of neural activity. The most commonly used linear method for dimensionality reduction is Principal Component Analysis (PCA), based on identifying mutually orthogonal directions in the empirical neural space of recorded activity; these directions are monotonically associated with the largest data variance. PCA provides a hierarchical description in which the data projected onto the manifold subtended by the principal components become closer and closer to the recorded data as the dimensionality of the linear manifold is increased towards the dimensionality of the empirical neural space. Although PCA provides a useful and systematic tool for variance-based dimensionality reduction, it does not specify how to uniquely identify the dimensionality of the manifold: the typical implementation requires the choice of an arbitrary variance threshold. Other PCA-based algorithms such as Participation Ratio (PR) [5, 18] and Parallel Analysis (PA) [31, 32] provide more principled prescriptions for linear dimensionality estimation, by incorporating criteria for determining an optimal number of leading principal components to use when constructing the low-dimensional manifold.

Linear dimensionality estimation algorithms may work well for linear datasets, but are likely to overestimate the dimensionality of a manifold arising from a nonlinear mapping between the low-and high-dimensional spaces [20, 21, 33, 34]. In contrast, nonlinear methods (e.g., Correlation Dimension [35-37], Levina-Bickel Maximum Likelihood Estimation [38], Two Nearest Neighbors [39], and Fisher Separability Analysis [40] may provide accurate dimensionality estimates for both linearly and nonlinearly embedded data.

Most dimensionality estimation methods have been tested in the absence of noise even though it is known that linear and nonlinear methods overestimate dimensionality when the data is noisy [20]. The robustness of dimensionality estimation algorithms to noise remains to be characterized.

The objective of this study was to characterize the accuracy of several dimensionality estimation algorithms when applied to high-dimensional recordings of neural activity. We evaluated previously proposed algorithms on synthetic datasets of known dimensionality to identify conditions under which each method succeeded and/or failed. Specifically, we evaluated how the algorithms handled the nature of the embedding (linear or nonlinear), the amount of noise added to the simulated neural data, and the amount of data available. We found increasing levels of noise to be a challenge for all tested algorithms. We therefore also evaluated different approaches for reducing noise prior to performing dimensionality estimation, including the “Joint Autoencoder”, a method we developed based on deep learning techniques. Together, our results allowed us to propose a methodological pipeline for estimating the intrinsic dimensionality of high-dimensional datasets of recorded neural activity.

## Methods

### Simulation of neural signals

We generated the synthetic data used to evaluate dimensionality estimation algorithms as follows. First, we created *d* signals by randomly sampling from an empirical distribution of firing rates that we obtained from multi-electrode array recordings of neural activity in the macaque primary motor cortex (M1) made while the subject was performing a center-out reaching task [41]. We verified that these randomly selected signals were uncorrelated. These signals provided a *d*-dimensional set used to construct synthetic high-dimensional data sets **(Fig 1)**. We allowed *d* to vary from 3 to 40. These signals were multiplied by a *d* x 96 mixing matrix *W* with entries that were randomly selected from a zero-mean Gaussian distribution with unit variance. This resulted in a 96-dimensional data set *X*. The activity in each of the *N*=96 simulated channels was scaled to the range from zero to one to compensate for variability in firing rates across neurons and across time. A nonlinear embedding was implemented by processing each simulated channel *X* with an exponential activation function:

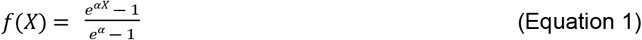

**Fig 1:**
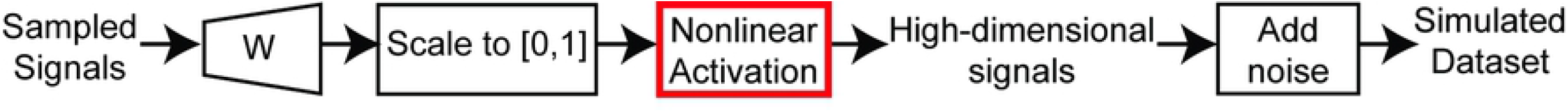
Generation of simulated datasets. First, representative neural signals were obtained by randomly sampling the firing rates of primary motor cortical recordings. The number of sampled signals determined the intrinsic dimensionality of the dataset. Then, the dimensionality of the sampled signals was increased through linear combinations by multiplying the signals with a weight matrix *W*. The entries of *W* were sampled from a zero-mean Gaussian distribution with unit variance. Then, the signals were then scaled to the [0,1] range by dividing them by their maximum value. This procedure yielded noise-free, linear datasets. In nonlinear simulations only, the signals were then activated nonlinearly through the exponential function in Equation 1 (red box in diagram). In noisy simulations, zero-mean Gaussian noise with variance specified by the predetermined signal-to-noise ratio was added to the signals. This procedure yielded linear or nonlinear, noisy datasets with known signal-to-noise ratio.

We chose this exponential activating function to control the degree of nonlinearity by varying the parameter *α*, and to ensure that the range of the nonlinearly embedded synthetic data remained between zero and one. Finally, we added independent Gaussian noise to each of the channels in *X*, to generate signals with known signal-to-noise ratio. This procedure generated datasets of known dimensionality, embedding type (linear/nonlinear), and signal-to-noise ratio.

### Dimensionality estimation algorithms

We evaluated two classes of dimensionality estimation algorithms, those that assumed a linear embedding and those that also allowed for a nonlinear embedding.

#### Linear algorithms

Linear algorithms map high-dimensional data to a lower dimensional, linear subspace. Principal Component Analysis (PCA) is often used for linear dimensionality estimation in neuroscience [2, 4, 7, 41-43]. All the linear algorithms that we tested (summarized below) are based on PCA but use different criteria for dimensionality estimation.

#### Principal Component Analysis with a variance cutoff

PCA creates a low-dimensional representation of the data by sequentially finding orthogonal directions that explain the most remaining variance. Unit vectors that identify those directions, the PCA eigenvectors {*ν_i_*}, provide an orthonormal basis for the *N*-dimensional data space. The eigenvectors are labeled in decreasing order of the variance associated with each direction, the eigenvalues {*λ_i_*}. The simplest way to use PCA for dimensionality estimation is to find the number of principal components required to reach a predetermined threshold of cumulative variance. The selection of a variance threshold can be rather arbitrary, and a range of thresholds have been used in the literature. In this study, we used a threshold of 90%, which yielded accurate estimates of dimensionality for the noise-free linear datasets.

#### Participation Ratio (PR)

This approach provides a principled way of finding a variance threshold when the ground truth is not known [5, 18]. PR uses a simple formula based on the eigenvalues:

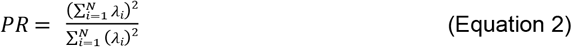

If the leading eigenvalue carries all the variance (*λi* ≠ 0 for *i = 1* and *λi* = 0 for all *i* ≥ 2), then PR = 1. At the other extreme, if all eigenvalues are equal, the variance is spread evenly across all the dimensions, and PR = *N*. The actual value of PR interpolates between these two extreme conditions to estimate the intrinsic dimensionality, and thus the number of principal components to be kept [5].

#### Parallel Analysis (PA)

Much like the Participation Ratio, Parallel Analysis is a principled approach to finding a variance threshold [31, 32]. Parallel Analysis generates null distributions for the eigenvalues by repeatedly shuffling each dimension of the data separately. The shuffling step ensures that the correlations remaining across the different dimensions of the data are due to chance. The eigenvalues that exceed the 95^th^ percentile of the null distribution are identified as significant, and their number is the number of dimensions to be kept. Although this method has not been directly applied to neural data, similar approaches based on finding null distributions of eigenvalues have been used for neural dimensionality estimation [44].

#### Nonlinear algorithms

Nonlinear algorithms can in principle estimate the dimensionality of either linearly or nonlinearly embedded data. Unlike the linear algorithms we tested, the nonlinear algorithms need not rely on a global model for the probability distribution from which the data are assumed to be drawn (in the case of PCA, the model is a multivariate Gaussian distribution). Instead, many nonlinear algorithms estimate intrinsic dimensionality directly from local geometric properties of the data. Common local properties include distance and separability of each data point relative to its neighbors. Although nonlinear algorithms are not yet commonly used in neuroscience, they have been used to estimate dimensionality in several other fields that produce high-dimensional datasets [45].

#### Correlation Dimension (CD)

Correlation Dimension estimates dimensionality by calculating how the number of data samples that fall within a hypersphere change as a function of its radius. This method, originally developed in 1983 [35], has benefitted from recent efforts to improve computational speed and accuracy [36, 37]. Although there are only a few applications of Correlation Dimension analysis to neural data [46, 47], it is widely used in other disciplines [36].

#### Levina-Bickel Maximum Likelihood Estimation (LBMLE)

The Levina-Bickel Maximum Likelihood Estimation method [38] is an extension of Correlation Dimension that uses a maximum likelihood approach to estimate distances between data points. This method has been successfully applied to some of the benchmark datasets used in machine learning, such as the Faces [33] and Hands datasets [48].

#### Two Nearest Neighbors (TNN)

The Two Nearest Neighbors method also uses the distance between data points to estimate dimensionality [39]. However, unlike Levina-Bickel Maximum Likelihood Estimation, it considers only the first and second neighbors of each point. The ratio of the cumulative distribution of second-neighbor to first-neighbor distances is a function of data dimensionality. By focusing on shorter distances, the method avoids unwanted effects resulting from density changes across the manifold. This method has been successfully applied to synthetic datasets of hyperspheres with known dimensionality [39], and to real-world datasets including molecular simulations [49] and images of hand-written digits [33].

#### Fisher Separability Analysis (FSA)

High-dimensional datasets exhibit simple geometric properties such as the likely orthogonality of two randomly picked directions. These properties have recently been characterized as the *blessings of dimensionality* [50], in contrast to the well-known concept of the *curse of dimensionality*. A useful example is the increasing ease with which a hyperplane can separate any given sample in a dataset from all other samples as the dimensionality of the dataset increases. Fisher separability is a computationally efficient, simple, and robust method to assess such separability [51, 52]. Dimensionality can be estimated in terms of the probability that a point in the dataset is Fisher separable from the remaining points [40]. The probability distribution of Fisher separability allows the dimensionality of both linear and nonlinear manifolds to be estimated. This method has been applied to study the mutation profiles of the genes resulting in tumors as a means to evaluate therapeutic approaches [53].

### Denoising algorithms

Noise that is uncorrelated across channels will lead to dimensionality estimates that approach the number of channels as the level of noise increases. To mitigate this overestimation problem, we implemented two approaches to denoise neural data. Both rely on an initial estimate of an upper bound dimensionality *D*, for which we used Parallel Analysis. To quantify the performance of the denoising algorithms, we reported variance accounted for (VAF) between the denoised signals and the noise-free signals.

#### PCA denoising

The linear approach to denoising was based on PCA. Once the value of *D* was determined, we used the *D* leading principal components to reconstruct the original data, under the assumption that most of the noise was relegated to the discarded, low-variance principal components.

#### Joint Autoencoder denoising

We also used a neural network for denoising **(Fig 2).** We divided the 96-dimensional simulated dataset *X* into two 48-dimensional partitions: *X*_1_ and *X*_2_. These partitions were each mapped by the compressive half of an autoencoder to compressed subspaces *Z*_1_ and *Z*_2_ respectively, each of dimension *D* < 48. These compressed subspaces were used to obtain reconstructed versions of *X*_1_ and *X*_2_, denoted 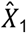 and 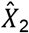, using the expansive halves of the corresponding autoencoders. The cost function *C* for the Joint Autoencoder network not only minimized the reconstruction error for *X*_1_ and *X*_2_, but also the difference between *Z*_1_ and *Z*_2_:

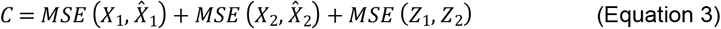

**Fig 2.**
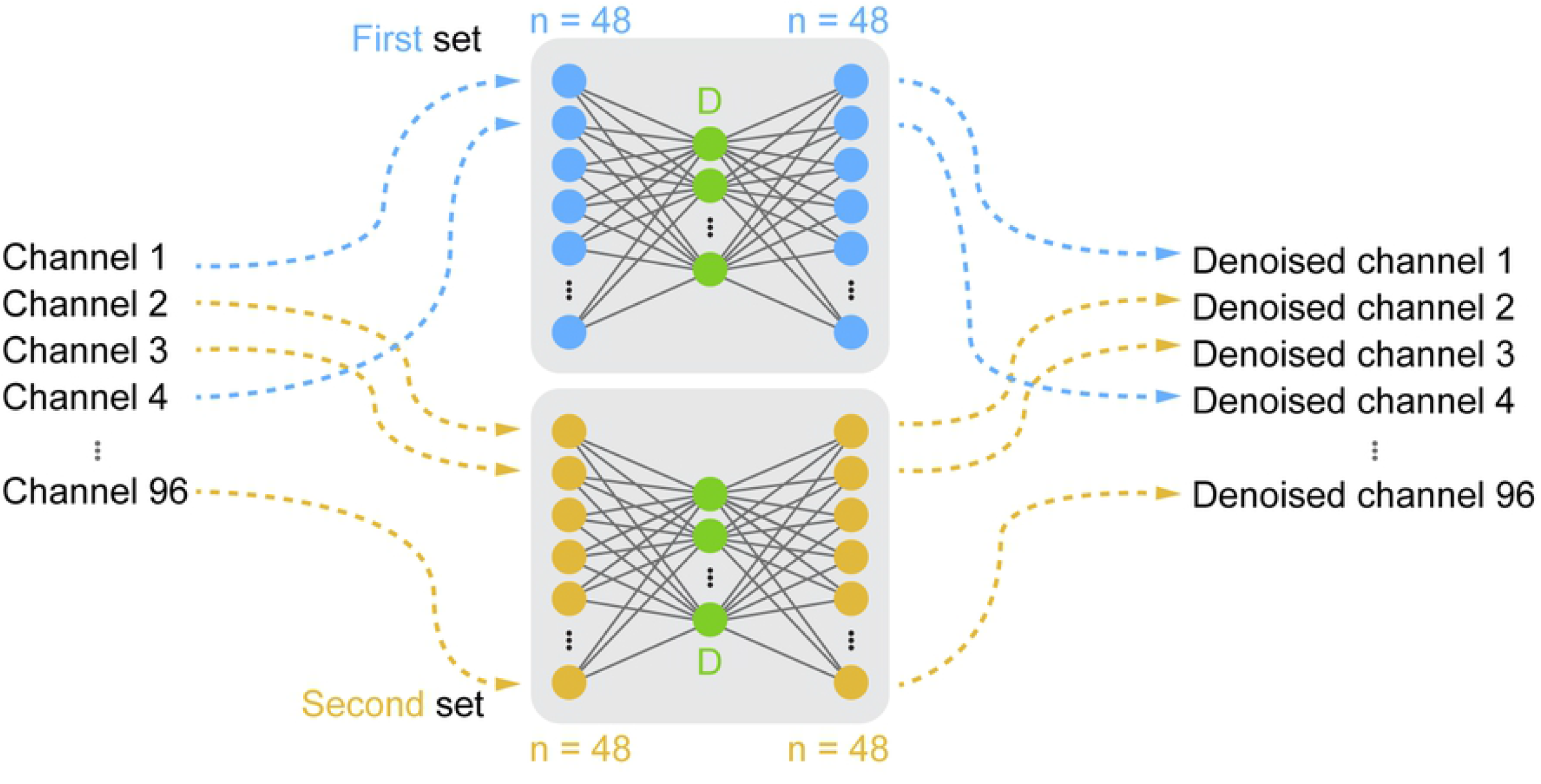
Architecture of the Joint Autoencoder. Channels of the 96-dimensional simulated datasets were randomly partitioned into two sets of signals (blue and yellow). Each 48-dimensional set was reconstructed through a D-dimensional subspace (green). The reconstructed outputs of the networks were the denoised channels.

This design assumes that each of the partitions *X*_1_ and *X*_2_ contains the information necessary to robustly identify the underlying *D*-dimensional signals *Z*_1_ and *Z*_2_, but not the independent noise components that will differ between the two partitions. We trained the Joint Autoencoder using the ADAM optimizer with a learning rate *η* = 0.001 and dropout regularization on the input layer with *p* = 0.05. The use of Rectified Linear Unit activation functions in all layers ensured that the autoencoder network would both operate on and output non-negative signals while allowing for nonlinear embeddings.

### Ethics statement

All surgical and experimental procedures that yielded the multi-electrode array recordings from non-human primates [41], which formed the basis of our simulated neural signals, were approved by Institutional Animal Care and Use Committee (IACUC) of Northwestern University. The subject was monitored daily. The subject’s diet consisted of standard laboratory animal diet, fresh fruits, and vegetables, and was provided with access to various types of enrichment.

### Statistical analyses

We used Monte Carlo simulations to generate up to 10 replications of synthetic data sets, each corresponding to microelectrode array recording data from an experimental session. We noted the number of replications (n) in the figure captions where applicable. Our choice of the number of replications is reasonable compared to the number of experimental sessions that we would expect to see in experiments with monkeys [41, 54, 55]. The simulations differed by their random number generator seed, which dictated the pseudorandom sampling procedures required for generating the signals. There were three sampling steps in our simulations **(Fig 1)**. First was the creation of the low-dimensional basis signals, which were sampled from an empirical firing rate distribution. The second was the entries of the mixing matrix *W*, which were sampled from a zero-mean Gaussian distribution with unit variance. The third was the additive noise, sampled from a zero-mean Gaussian distribution with variance determined by the signal-to-noise ratio. We used bootstrapping with 10,000 iterations to compute the statistic of interest and computed its confidence interval using α = 0.05. We used Bonferroni correction for multiple comparisons.

## Results

Despite the large number of available algorithms for dimensionality estimation, there has been no systematic study of how well-suited they are for the analysis of neural data. Here we test several representative algorithms on synthetic datasets for which the intrinsic dimensionality is known, to assess their ability to estimate the true dimensionality of the data across a range of simulated conditions relevant to neuroscience. These assessments resulted in a recommended procedural pipeline for estimating the intrinsic dimensionality of a set of neural recordings.

### Dimensionality of noise-free datasets

We first considered the simplest case: how accurately can we determine the dimensionality of linearly embedded, noise-free datasets? To answer this question, we applied the six selected algorithms to datasets with dimensionality *d* = 6. We focused on *d* = 6 as this was the dimensionality estimate of actual multi-electrode array recordings found when using the methods investigated here. In this scenario, all tested linear and nonlinear algorithms estimated the true dimensionality accurately **(Fig 3)**. Under noise-free conditions, the nonlinear algorithms were as accurate as the linear ones on linearly embedded datasets.

**Fig 3.**
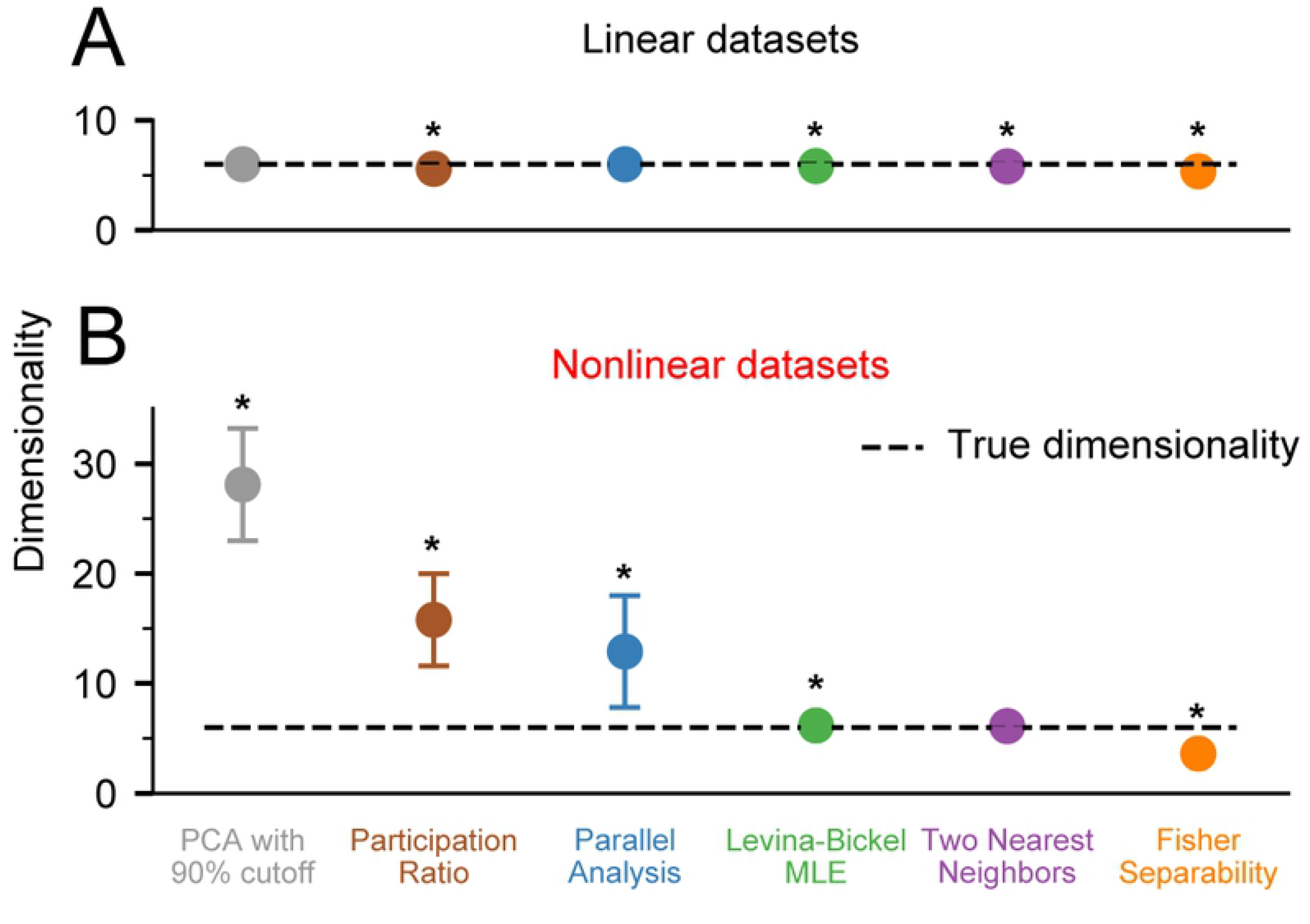
Dimensionality of noise free datasets. A) We applied PCA with 90% variance cutoff (PCA90, gray), Participation Ratio (PR, brown), Parallel Analysis (PA, blue), Levina-Bickel Maximum Likelihood Estimation (LBMLE, green), Two Nearest Neighbors (TNN, purple), and Fisher Separability Analysis (FSA, orange) to linearly embedded, *d* = 6 datasets (n=10). B) Same as in A, but for nonlinearly embedded datasets. Circles indicate the mean and error bars indicate the standard deviation of the dimensionality estimates. Asterisks indicate significant difference of the mean from the true dimensionality of 6 at (bootstrapped confidence intervals do not overlap 6 at the significance level of α=0.05.

Next, we evaluated all algorithms on nonlinearly embedded noise-free datasets, also for *d* = 6. Nonlinearities were introduced as in Equation 1, using α = 16. In this case, the three linear algorithms dramatically overestimated the true dimensionality, with errors reaching more than 400% of the true value **(Fig 3B)**. In contrast, the nonlinear algorithms performed well; the Levina-Bickel Maximum Likelihood Estimation and the Two Nearest Neighbors methods were more accurate than Fisher Separability Analysis, which slightly underestimated the true dimensionality.

Because of the superior accuracy of Levina-Bickel Maximum Likelihood Estimation and Two Nearest Neighbors, we focused on these two methods for the remainder of the nonlinear analyses. We also retained Parallel Analysis as a benchmark for some of the analyses, as it was the most accurate linear method for estimating the dimensionality of nonlinearly embedded data.

### Effect of true dimensionality on algorithm accuracy

We next evaluated how the true intrinsic dimensionality of the noise-free data influenced algorithm accuracy. Can any intrinsic dimensionality be reliably estimated? We found that the answer is no: the accuracy of all algorithms suffered when the intrinsic dimensionality of the synthetic data was too high. Parallel Analysis was accurate on linear datasets with *d* < 20, but inaccurate on nonlinear datasets of all dimensions, as expected **(Fig 4)**. Below about *d* = 6, Levina-Bickel Maximum Likelihood Estimation and Two-Nearest Neighbors were accurate on both linear and nonlinear datasets. However, Levina-Bickel Maximum Likelihood Estimation began to underestimate the dimensionality of both linearly embedded **(Fig 4A)** and nonlinearly embedded **(Fig 4B)** datasets for *d* > 6. This underestimation increased with increasing *d*. For nonlinear datasets, the estimate saturated at *d* = 13, where underestimation began to get much worse. These results revealed that the intrinsic dimensionality of nonlinearly embedded datasets is hard to estimate reliably when it is large.

**Fig 4.**
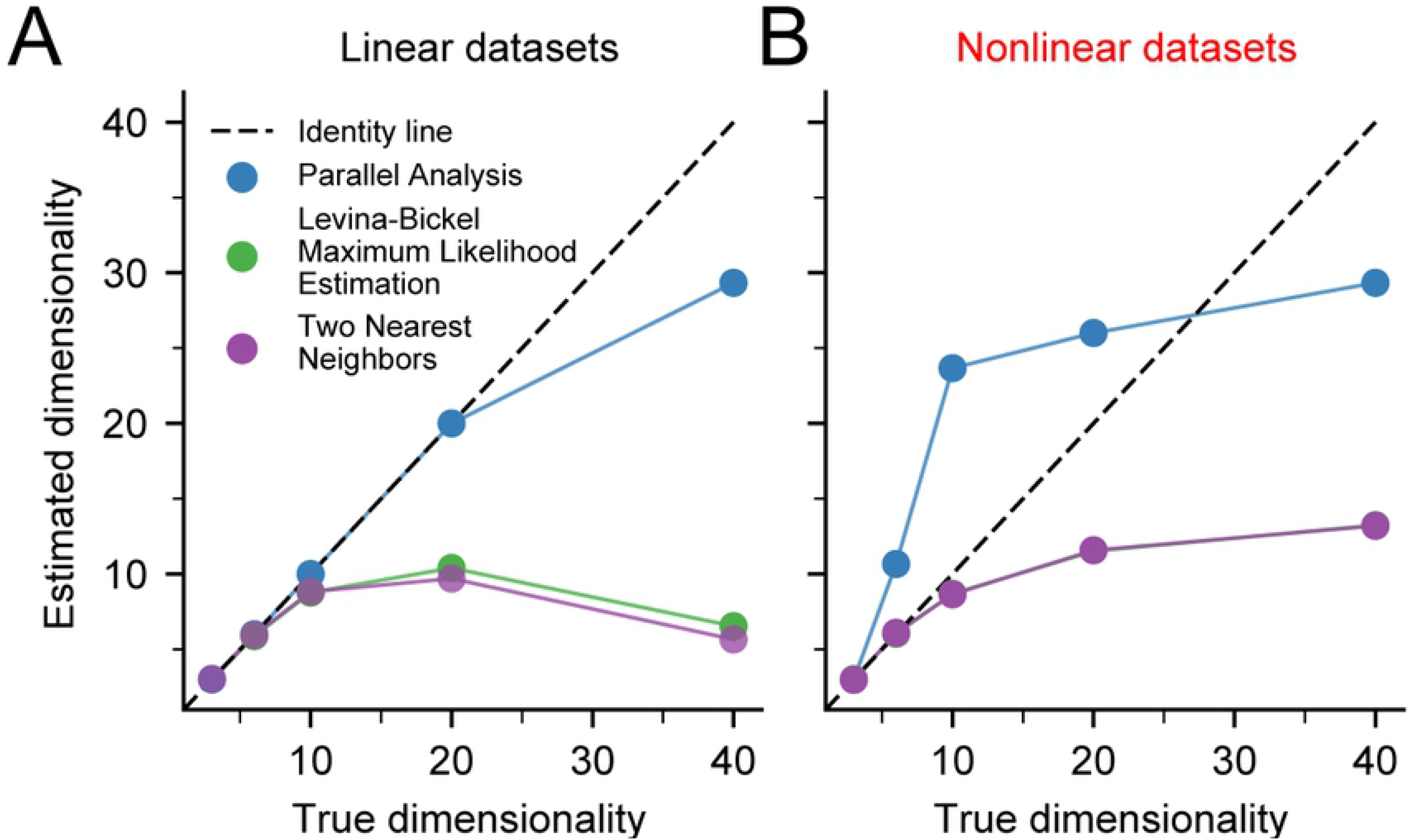
Effect of increasing true dimensionality on dimensionality estimates. A) The dimensionality of noise free, linear datasets (n=3) was assessed using Parallel Analysis (PA), Levina-Bickel Maximum Likelihood Estimation (LBMLE), and Two Nearest Neighbors (TNN). Dashed line indicates the identity line. B) Same as A, but for nonlinear datasets. The curve for TNN precisely overlays that of LBMLE, causing it to be obscured.

### Effect of the level of nonlinearity

We next evaluated how the degree of nonlinearity influenced the accuracy of the dimensionality estimation algorithms. We controlled the degree of nonlinearity by varying the parameter α in Equation 1; this parameter controls the slope of the exponential activation function used to generate the nonlinearly embedded datasets. We found that both Levina-Bickel Maximum Likelihood Estimation and Two Nearest Neighbors provided accurate dimensionality estimates for all tested levels of nonlinearity (**Fig 5**). Surprisingly, even Parallel Analysis was accurate at levels of nonlinearity around *α* ≈ 8, where it started to overestimate the intrinsic dimensionality. These results revealed that Levina-Bickel Maximum Likelihood Estimation and Two Nearest Neighbors provide accurate dimensionality estimates for wide levels of nonlinearity, whereas Parallel Analysis is accurate only for low levels of nonlinearity.

**Fig 5.**
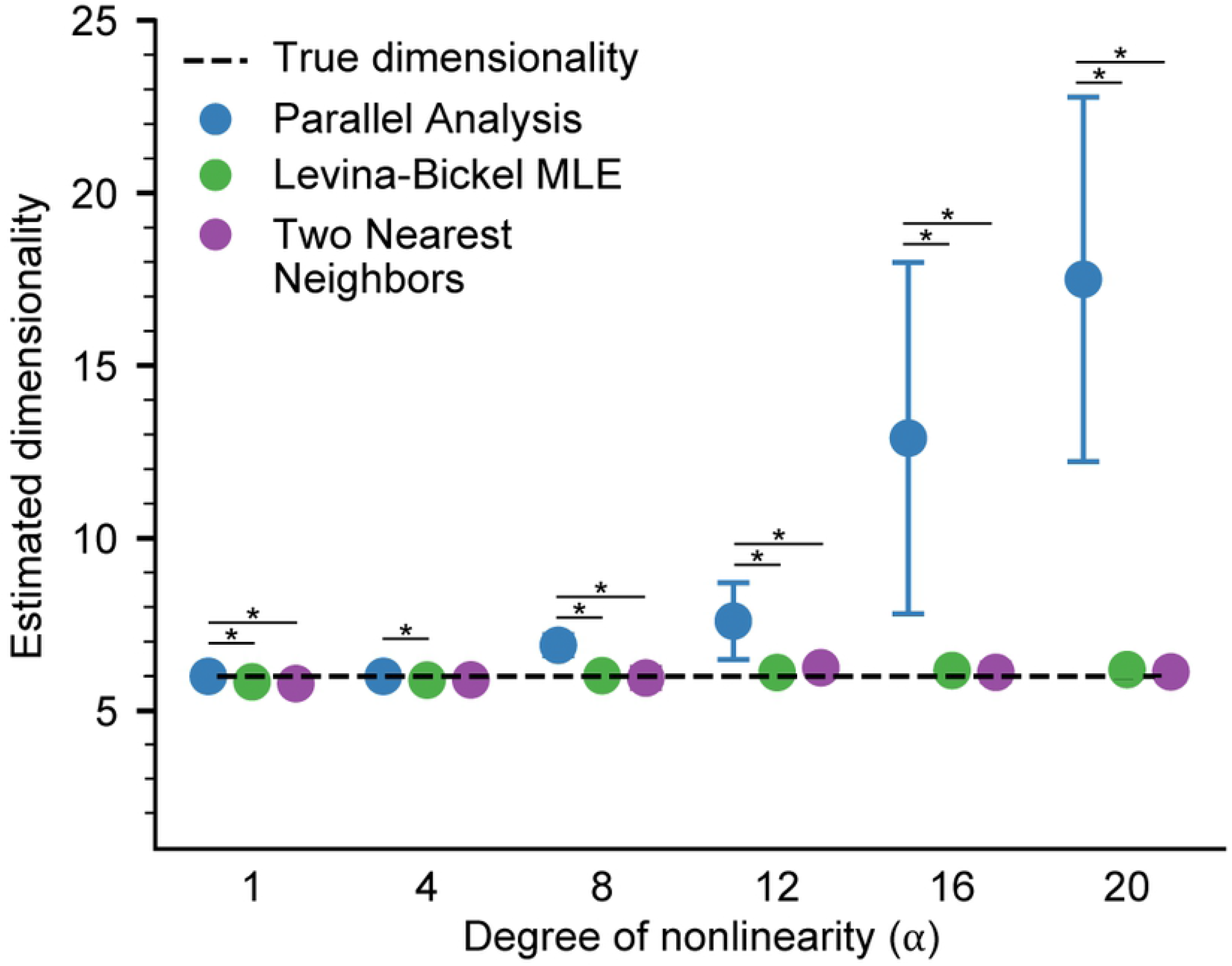
Effect of changing the degree of nonlinearity. Dimensionality of nonlinear datasets (n=10) with varying levels of nonlinearity, controlled by the α parameter (See Methods), was assessed using Parallel Analysis (PA), Levina-Bickel Maximum Likelihood Estimation (LBMLE), and Two Nearest Neighbors (TNN). Circles indicate the mean and error bars indicate the standard deviation of the dimensionality estimates. Asterisks indicate significant difference between mean values (bootstrapped confidence intervals do not overlap 0 at the significance level of α=0.05/3, Bonferroni corrected for multiple comparisons).

### Amount of data required for estimating dimensionality

Ideally, algorithms would require only small amounts of data, so that the intrinsic dimensionality could be estimated even during transient behaviors and for a small number of recording channels. We thus evaluated the amount of data required to estimate dimensionality of datasets with *d* = 6, by varying both the duration of the recordings and the number of recording channels.

On linear datasets, the accuracy of Parallel Analysis depended only on the number of channels: the algorithm was accurate if 20 or more channels were available (**Fig 6A**). In contrast, the accuracy of both Levina-Bickel Maximum Likelihood Estimation and Two Nearest Neighbors also depended on the duration of the data (**Fig 6B and C**). Around 30 seconds of data was sufficient for accurate estimates of intrinsic dimensionality using either of these two nonlinear methods.

**Fig 6.**
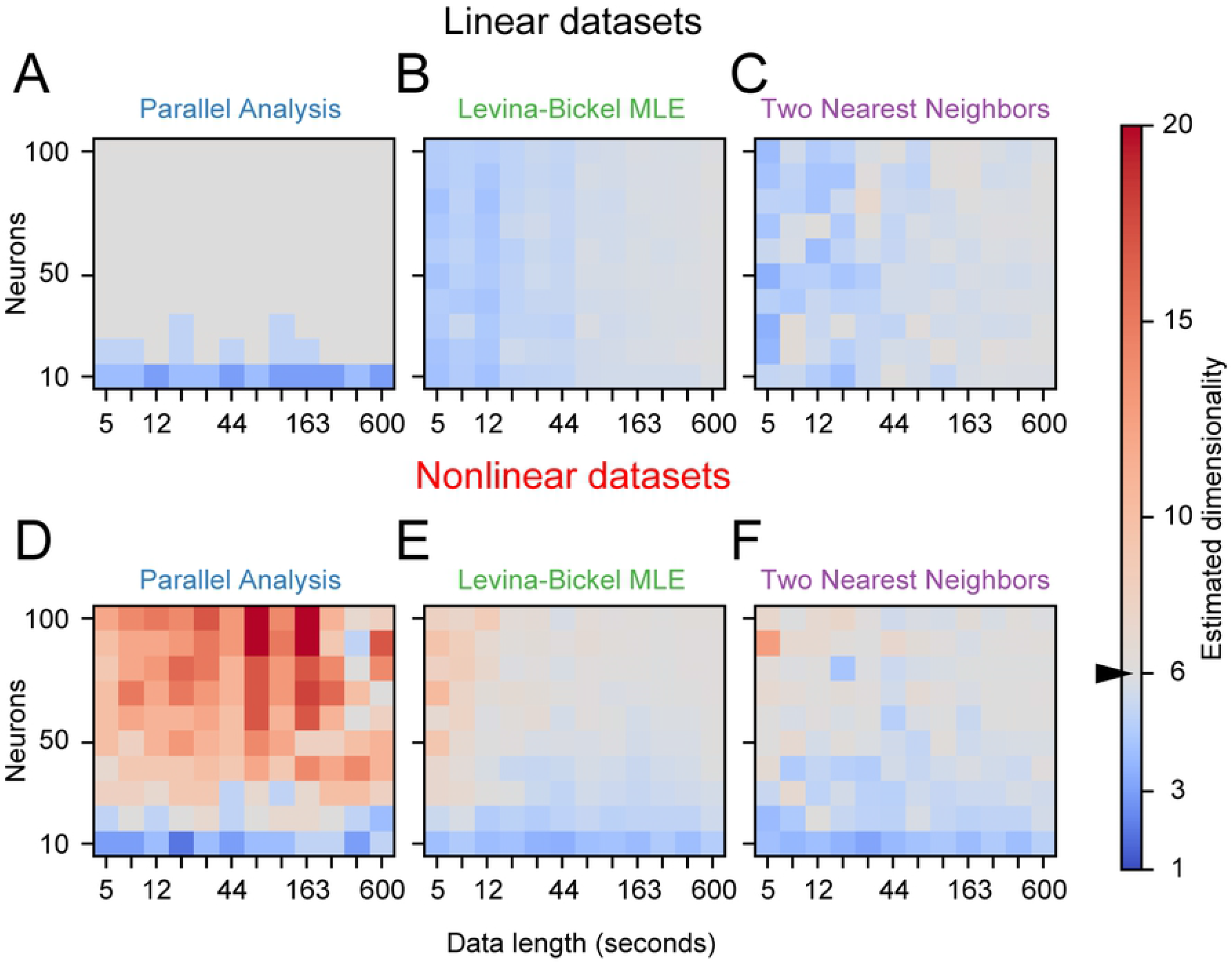
Amount of data required by dimensionality estimators. Amount of data required by A) Parallel Analysis (PA), B) Levina-Bickel Maximum Likelihood Estimation (LBMLE), and C) Two Nearest Neighbors (TNN) on linear datasets. Data length is logarithmically scaled between 5 seconds and 600 seconds. Correct dimensionality *d* = 6 is shown in gray. Warm colors indicate overestimation and cold colors indicate underestimation of dimensionality. D, E, and F) Same as A, B, and C, respectively, but for nonlinear datasets.

As expected for highly nonlinear datasets (*α* = 16 *d*, = 6), Parallel Analysis was not accurate (**Fig 6D**) regardless of the amount of data. Both Levina-Bickel Maximum Likelihood Estimation and Two Nearest Neighbors were accurate provided that data from more than 50 channels were available (**Fig 6E and F**). Furthermore, while Levina-Bickel Maximum Likelihood estimation required around 30 seconds of data for accurate dimensionality estimates, Two Nearest Neighbors required more than one minute. These results would also depend on the actual dimensionality *d* of the tested signals; here we focused on *d* = 6.

### Evaluating and reducing the effects of noise

Any experiment will include some amount of noise in the recorded signals. As expected, all tested algorithms overestimated intrinsic dimensionality in the presence of noise (**Fig 7**). For any given noise level, estimation errors for the linear datasets **(Fig 7A)** were a bit smaller than those for the nonlinear datasets **(Fig 7B)**. Adding noise with a power of only 1% of that of the signal (SNR = 20 dB) caused Levina-Bickel Maximum Likelihood Estimation and Two Nearest Neighbors to overestimate the dimensionality of the nonlinear data by ~200% **(Fig 7B)**. PA yielded consistent overestimation errors across all nonzero levels of noise for both linear and nonlinear data.

**Fig 7.**
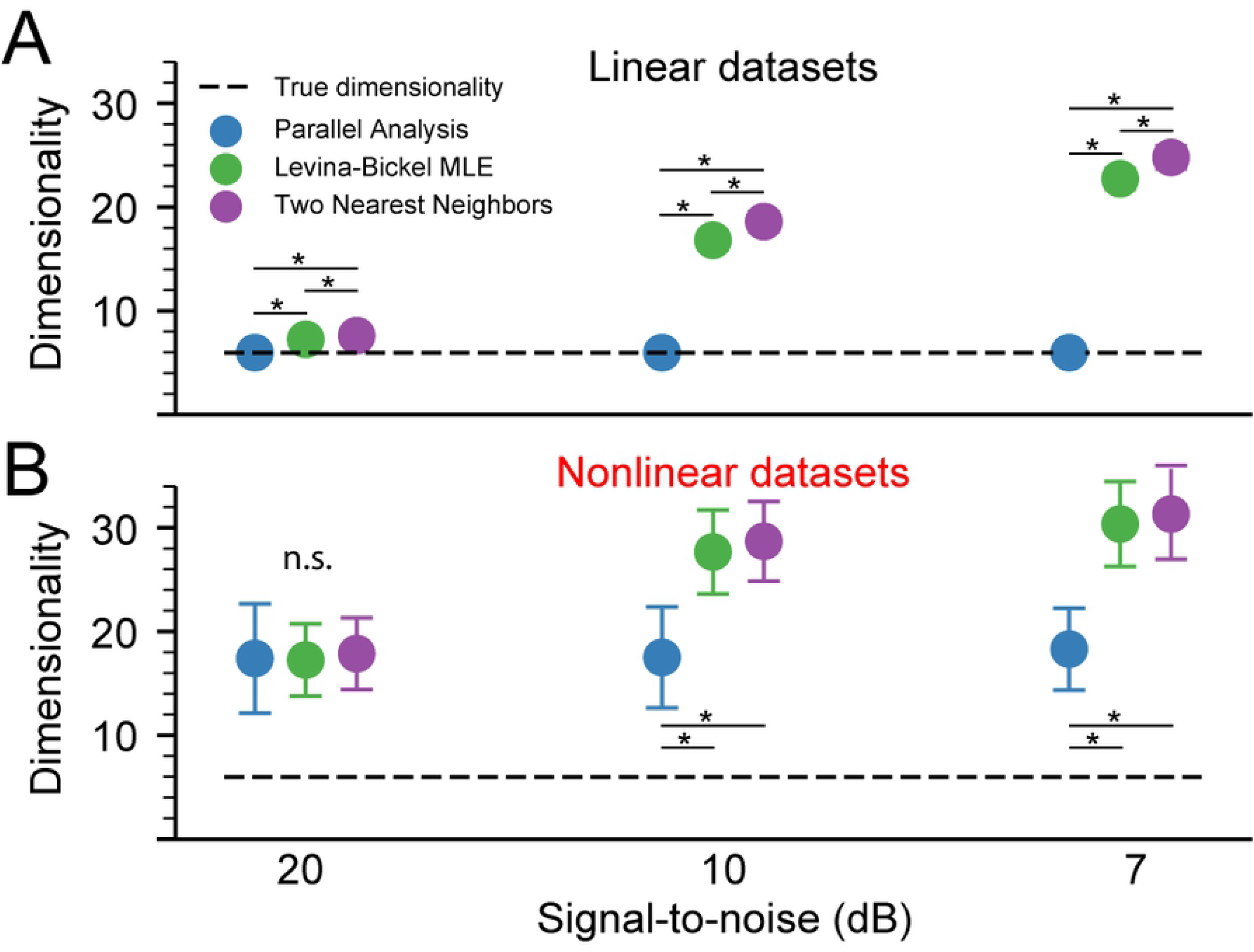
Effect of noise on dimensionality estimates. Estimated dimensionality of linear (A) and nonlinear (B) datasets (n=10) with 20 dB, 10 dB, and 7 dB signal-to-noise ratio was assessed using Parallel Analysis (PA), Levina-Bickel Maximum Likelihood Estimation (LBMLE), and Two Nearest Neighbors (TNN). Circles indicate the mean and error bars indicate the standard deviation of the dimensionality estimates. Asterisks indicate significant difference between mean values (bootstrapped confidence intervals do not overlap 0 at the significance level of α=0.05/3, Bonferroni corrected for multiple comparisons).

We evaluated two algorithms for mitigating the effects of noise prior to estimating dimensionality: a PCA-based linear method and a Joint Autoencoder nonlinear neural network (see Methods). Both methods were quite effective for denoising the linear datasets **(Fig 8A)**, with the PCA-based approach slightly better than the Joint Autoencoder at the higher noise levels. For linear datasets, dimensionality estimates following PCA-based denoising were highly accurate, yielding correct estimates of the true intrinsic dimension even for high-noise signals **(Fig 8B)**. The Joint Autoencoder was significantly more effective for denoising the nonlinear datasets **(Fig 8C)**. Joint Autoencoder denoising on nonlinear datasets resulted in dimensionality estimates that still increasingly overestimated with increasing noise, but at a much slower rate than without denoising **(Fig 8D)**. The highest noise level we tested (20%; SNR = 7 dB) caused the dimensionality to be overestimated by about 100%.

**Fig 8.**
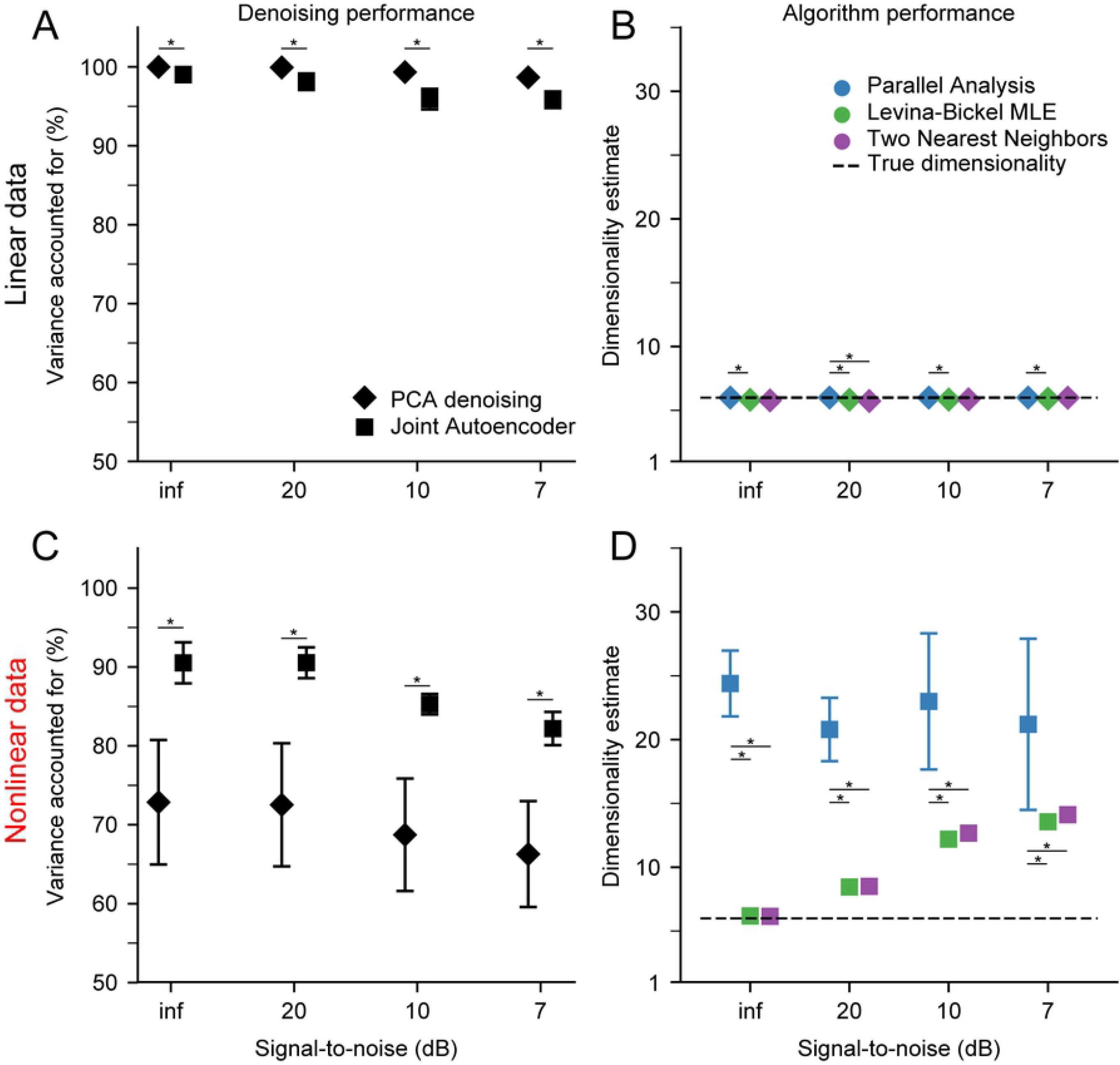
Performance of PCA and Joint Autoencoder (JAE) denoising algorithms. A) PCA and JAE denoising applied to linear datasets (n=10) with varying signal-to-noise ratio. Symbols indicate the mean and error bars indicate the standard deviation of the Variance accounted for between noise-free and denoised signals. Asterisks indicate significant difference between mean values (bootstrapped confidence intervals do not overlap 0 at the significance level of α=0.05). B) Dimensionality estimation on linear datasets after PCA denoising. Dimensionality was estimated using Parallel Analysis (PA), Levina-Bickel Maximum Likelihood Estimation (LBMLE), and Two Nearest Neighbors (TNN). Symbols indicate the mean and error bars indicate the standard deviation of the dimensionality estimates. Asterisks indicate significant difference between mean values (bootstrapped confidence intervals do not overlap 0 at the significance level of α=0.05/3, Bonferroni corrected for multiple comparisons). C) Same as in A, but for nonlinear datasets. D) Same as in B, but for nonlinear datasets after JAE denoising.

## Discussion

This study evaluated techniques for estimating the intrinsic dimensionality of high-dimensional neural recordings. We considered representative linear and nonlinear algorithms, testing their performance on synthetic datasets that captured properties of neural recordings likely to affect dimensionality estimation. The tested datasets had known intrinsic dimensionality, known levels of noise, and embeddings that were either linear or nonlinear. Our results demonstrated that none of the tested algorithms work for all possible scenarios, but they yielded important insights for when estimates of intrinsic dimensionality are likely to be valid and when they are not. As expected, we found that linear estimation methods are generally not as accurate as nonlinear methods when the mapping between the low-dimensional latent space and the high-dimensional space of neural recordings is nonlinear. Surprisingly, the linear method Parallel Analysis estimated the dimensionality of mildly nonlinear datasets well though it failed for more highly nonlinear embeddings. In contrast, the nonlinear methods worked well on both linear and highly nonlinear datasets but failed once the intrinsic dimensionality of the data became too high.

Noise was a challenge for all methods, causing dimensionality to be overestimated even for signal-to-noise ratios as low as 20 dB (1% noise variance). We presented two approaches for denoising the data so as to improve the accuracy of the dimensionality estimation. These were a linear PCA-based approach and a novel nonlinear, deep learning approach that we call the Joint Autoencoder. Both denoising approaches attempted to remove signal components that were not shared across the data channels. To achieve this, the PCA-based approach simply removed Principal Components with low variance, whereas the Joint Autoencoder identified an underlying manifold that was common to two randomly sampled sets of channels. Both approaches relied on a linear, upper-bound estimate of the intrinsic dimensionality. Denoising by either method substantially improved subsequent dimensionality estimation, but the Joint Autoencoder was substantially more effective in denoising nonlinear datasets. In the linear case, dimensionality estimates using Parallel Analysis, Levina-Bickel Maximum Likelihood Estimation, and Two Nearest Neighbors were accurate after PCA-denoising. In the nonlinear case, dimensionality estimates using the same three methods were similarly accurate after JAE-denoising.

### Implications for evaluation of experimental recordings

Due to its computational efficiency and ease of interpretation, most studies have used PCA with an arbitrary variance cutoff to estimate the dimensionality of M1 neural recordings [4, 17, 41-43]. While we have shown that some of the linear methods can be quite effective, simply eliminating non leading PCs based on a cumulative variance cutoff was the least accurate of the algorithms that we tested. Parallel Analysis, the most accurate linear method, performed as well or even better than some of the more advanced and computationally demanding nonlinear methods. Therefore, PA should suffice as a quick and effective approach to estimating dimensionality, even for mildly noisy and nonlinear datasets.

Despite the simplicity of linear algorithms, estimating dimensionality of nonlinear manifolds requires nonlinear algorithms. There is some evidence that neural manifolds may be nonlinear. Recent studies have shown that nonlinear methods for “decoding” behavioral parameters from M1 neural manifolds are superior to linear methods [56-59]. This suggests that the underlying neural manifold representing motor intent may be nonlinear, and that linear dimensionality estimation methods may be inadequate when estimating the intrinsic dimensionality of primary motor cortical recordings. Studies that investigated the dimensionality of M1 using linear methods most likely overestimated its true intrinsic dimensionality.

Nonlinear algorithms were more accurate than linear methods for nonlinear datasets of dimensionality below 10. However, nonlinear methods underestimated dimensionalities above 10. This is a critical concern for experimental recordings, since a low dimensionality estimate from a nonlinear method might be inaccurate if the true dimensionality were large. Multiple studies using linear methods have reported an estimated dimensionality of M1 of around 10 for simple, well-practiced behaviors [5, 43, 55]. Our results show that linear methods provide an upper bound to the estimate of intrinsic dimensionality as long as the true dimensionality of the data is below 20. If the intrinsic dimensionality of M1 is substantially higher for more dexterous use of arm and hand than for the scenarios that have typically been studied, the nonlinear methods investigated here may underestimate it.

One method for addressing this concern would be to use nonlinear methods to reduce the dimensionality of a dataset to that of its nonlinear dimensionality estimate, and then to assess the amount of variance that the nonlinear low-dimensional representation captures. If the resulting variance accounted for (VAF) is high, the data may be truly nonlinearly low dimensional. If, on the other hand, the VAF is low, the true intrinsic dimensionality could be higher than estimated. For the latter case, a practical approach would be to report only the linear dimensionality estimate and emphasize that it only provides an upper bound to the true dimensionality.

We currently lack techniques for reliably assessing datasets with high intrinsic dimensionality, at least when considering practical situations with limited data. There have been some theoretical studies of the amount of data needed for accurate estimation of dimensionality [60, 61]. Correlation Dimension, the method on which many nonlinear algorithms are based, requires that the number of data samples be on the order of 10^d/2^ [29]. The amount of data can be increased by either recording from more channels or for a longer duration. One study that investigated the dimensionality of the primary visual cortex (V1) found the eigenvalue spectrum of the neural signals obtained from approximately thousand neurons to decay as a power law [62]. Their finding would not have been possible had they recorded from fewer neurons, which would have prevented them from observing the long tail of the eigenvalue distribution. One interpretation of this finding is that the linear dimensionality is arbitrarily large. However, an alternative interpretation is the existence of an extremely nonlinear manifold embedded within the neural space investigated in that study.

The stochastic nature of neural firing and the noise associated with experimental measurements will also cause the intrinsic dimensionality to be overestimated. The two denoising approaches that we presented are simple and effective. Depending on the assumptions about the underlying structure of firing patterns, alternative denoising approaches may be useful. For example, if the temporal relationship between the firing patterns of the population neural activity is of interest, one could use denoising methods that explicitly attempt to model these dynamics, such as Latent Factor Analysis through Dynamical Systems (LFADS), prior to estimating the dimensionality [57].

### Limitations of the study

While we tried to replicate essential features of experimental data, there are certain characteristics that we did not try to model in our simulations. For example, we only considered additive Gaussian isotropic noise, for simplicity. Experimental recordings might include non-additive, non-isotropic, or non-Gaussian noise. In such cases, PCA may not be an appropriate approach to denoising, even for linearly embedded data. Methods such as factor analysis or extensions such as Gaussian-Process Factor Analysis [63], and preprocessing steps such as square-root transforms or pre-whitening could be used instead.

We scaled the firing rates of each channel to be in the [0,1] range. This procedure does not reflect experimental neural firing data, since the range of neural firing can differ significantly even across neurons of the same type. The arbitrary scaling of firing rates provided a simple means for the nonlinear datasets to have the same range as their linear counterparts, as the activation function that we used mapped the [0,1] range onto itself.

### Recommended analysis pipeline

Based on our results, we recommend the following approach for estimating the dimensionality of neural recordings. First, obtain an upper-bound estimate *D* of the intrinsic dimensionality of the data. We found that Parallel Analysis works well for this purpose, being both computationally efficient and the most accurate linear method in our tests. Next, the signals should be denoised. Our denoising approach worked by projecting the neural signals into a subspace of dimensionality *D* equal to the upper-bound dimensionality estimate, and then reconstructing them based on these projections. A PCA based reconstruction is easy to implement and interpret and may be preferable if computational efficiency is important. A nonlinear denoising algorithm, such as the Joint Autoencoder we proposed, should also be used to assess the nonlinearity of the manifold. The usefulness of the denoising step was quantified through the variance accounted for (VAF) between the reconstructed signals, assumed to be denoised, and the noise-free synthetic signals before noise was added to them. Our results showed that for nonlinear datasets this VAF was higher for the Joint Autoencoder than it was for PCA. However, this VAF cannot be computed for experimental data, for which we do not have access to the noise-free signals. In this scenario, the reconstruction VAF between noisy inputs and the denoised reconstructed outputs may be useful for detecting nonlinear manifolds: a higher reconstruction VAF for Joint Autoencoder denoising than for PCA denoising would signal a nonlinear manifold. If the reconstruction VAF results prefer the Joint Autoencoder, this denoising method yields better denoised signals. Once the signals are denoised, and the linearity of the manifold is established, either a linear or nonlinear dimensionality estimation method should be used depending on the expected linearity of the manifold as determined by the comparative performance of the denoising algorithms. The most accurate linear method we tested was Parallel Analysis. Of the nonlinear methods, Levina-Bickel Maximum Likelihood Estimation and Two Nearest Neighbors were the most accurate; Levina-Bickel Maximum Likelihood Estimation required fewer samples.

## Conclusions

Estimating the dimensionality of neural data is challenging. In this study, we tested several available algorithms, and determined the conditions under which estimating dimensionality may be particularly difficult or even impractical. Noise is a confounding factor and must be eliminated prior to dimensionality estimation. Most existing studies have estimated intrinsic dimensionality using linear methods, as they are computationally efficient and easy to interpret. We showed that linear methods provide an upper-bound to the intrinsic dimensionality, and in cases of high noise, may even work better than nonlinear methods, although neither linear nor nonlinear methods will yield accurate estimates in this scenario. Nonlinear algorithms were more accurate for nonlinear datasets when noise was adequately removed. Finally, algorithms failed when the intrinsic dimensionality was high. It may be impractical or impossible to estimate the dimensionality of neural data when it is above ~20. However, estimation of the dimensionality of neural activity in the primary motor cortex may be possible, as many studies have reported its linear dimensionality to be within the practical limits for accurate estimation by the methods we tested.

## Acknowledgments

The authors would like to thank Juan Á. Gallego for collecting the experimental data which served as the basis for our simulations.

